# Toxoplasmicidal *in vitro* effect of dehydroepiandrosterone on Toxoplasma gondii extracellular tachizoytes

**DOI:** 10.1101/2020.03.23.004713

**Authors:** Saé Muñiz-Hernández, Carmen T. Gómez de León, Angélica Luna Nophal, Lenin Domínguez-Ramírez, Olga-Araceli Patrón-Soberano, Karen E Nava-Castro, Pedro Ostoa-Saloma, Jorge Morales-Montor

**Affiliations:** Laboratorio de Oncología Experimental, Subdirección de Investigación Básica, Instituto Nacional de Cancerología, Secretaria de Salud, Ciudad de México, 14080, México; Departamento de Inmunología, Instituto de Investigaciones Biomédicas, Universidad Nacional Autónoma de México, AP 70228, Ciudad de México, 04510, México; Departamento de Ciencias Químico-Biológicas, Escuela de Ciencias, Universidad de las Américas Puebla, Santa Catarina Mártir, Cholula, Puebla, 72810, México; División de Biología Molecular, Instituto Potosino de Investigación Científica y Tecnológica, Camino a la Presa San José 2055, Col. Lomas 4a. sección, San Luis Potosí, 78216, México; Laboratorio de Genotoxicología y Mutagénesis Ambientales, Departamento de Ciencias Ambientales, Centro de Ciencias de la Atmósfera, Universidad Nacional Autónoma de México. Ciudad de México, 04510, México

**Keywords:** Toxoplasmosis, *Toxoplasma gondii*, tachyzoite, dehydroepiandrosterone, in vitro, parasiticide effect, drug

## Abstract

Toxoplasmosis is a zoonotic disease caused by the apicomplexa protozoan parasite *Toxoplasma gondii*. This disease is a health burden, mainly in pregnant women and immunocompromised individuals, in whom they can cause death. Despite advances in the medical area, nowadays there are no new drugs to treat toxoplasmosis. The standard therapy to toxoplasmosis has not had progress for last seven decades; it is a combination of sulfadiazine-pyrimethamine (S-P); which is co-administered with folic acid due to the adverse effects of the drug. Several studies have shown that the conventional treatment has limited effectiveness and severe adverse effects. Thus, the search of better treatments with greater efficacy and without the adverse effects becomes relevant. In the current work we demonstrate for the first time the parasiticidal effect of dehydroepiandrosterone (DHEA), a steroid hormone produced by many mammals, on extracellular tachyzoites (the infective stage of *T. gondii*). In vitro treatment with DHEA reduces the viability of extracellular tachyzoites, and both the active and passive invasion processes. The ultrastructural analysis of treated parasites showed that DHEA alters the cytoskeleton structures, leading in the lost of the organelle structure and organization, as well as, the lost of the cellular shape. On a molecular level, we observed an important reduction of the expression of several proteins that are essential for the motility and virulence of parasites when they were exposed to DHEA. These results suggest that DHEA could be used as an alternative treatment against toxoplasmosis.

## Introduction

Toxoplasmosis is a zoonosis caused by the apicomplexa protozoan parasite, *Toxoplasma gondii*, which is able to infect all warm-blooded animals [1-2]. This is a worldwide disease with a prevalence average of 40 % [3]. Particularly, in Mexico the sero-prevalence goes from 40-70% depending on the region of the country [4-5]. *T gondii* infection can induce abortion, encephalitis, and in extremely cases, death. It is considered a major opportunistic pathogen in patients with AIDS [6-7].

Human toxoplasmosis presents two phases: the acute and the chronic. In the acute phase, parasite disseminate in the tachyzoite stage, the highly invasive and motile asexual form. In this stage, parasite is able to cross any biological barrier, included the placenta or the blood-brain barrier [8-11]. If the host is immunocompetent tachyzoites will eventually differentiate into bradyzoites, the low replication form, and will begin the tissue cyst formation [12]; this event defines the chronic infection, since tissue cysts can stay forever in the host without provoking any apparent pathology [13].

Tachyzoite stage has a characteristic half-moon shape and an approximate size of 5 to 10 μm [14], as all members of Apicomplexa family; its motility depends on actomyosin machinery that underlies the plasma membrane called glideosome [15]. *Toxoplasma* counts with three specialized secretory organelles with particular proteins, which are secreted in a controlled and specific manner during biological process, the micronemes (MIC protein), rhoptries (ROP proteins) and dense granules (GRA proteins) [16].

*Toxoplasma* tachyzoite can carry out two types of invasion, active or passive. In the active invasion *Toxoplasma* is the effector cell, and is the most important process due to at majorly of the cell in the individuals are infected by mean this process. Firstly, the tachyzoite must adhere to plasma membrane of the host cell; then glides propelled by the glideosome that links to the host cell membrane via MIC2/MAP2 complex. It has been described that parasite recognizes an unknown ligand of the host by its GPI-anchored surface antigens, known as SAGs. Then, micorneme protein AMA1 and RON proteins (RON2, RON4, RON5 and RON8) are secreted and a fusion of both plasma membranes, called moving junction (MJ) is established [13, 17-18]. Parasite twirls inside of the host cell at the same time that the PV is formed by the secretion of ROP and GRA proteins. Inside of the non-fusogenic PV, the parasite is replicated by endodiogeny, an asexual replication form, that from the boundaries of a mature mother parasite forms two daughters’ cells [19-20].

Passive invasion occurs in all phagocytic cells, these being the effector cells for the process. First, the parasite adheres to the plasma membrane of a phagocytic activated cell, surrounded by the plasmatic membrane elongations and internalized towards the cytoplasm in a phagocytic vacuole [13]. Once inside, the parasite evades the immune response transforming the phagocytic vacuole in to a parasitophorous vacuole (PV) via the phosphorylation of the host Immune-Related GTPases (IRGs) via a complex that includes ROP and GRA proteins. This prevents their oligomerization and recruitment to the PVM leading in the inhibition of the vacuole lysis and parasite clearance by macrophages then parasite is able to replicate [13, 21-23].

Conventional therapy against toxoplasmosis consists of a mixture of sulfadiazine-pyrimethamine that was established in the 50’s decade. Since then, minor advances have been made in the treatment of the zoonosis [24-26]. Although sulfadiazine – pyrimethamine are synergic it is known that they present severe side effects. Since pyrimetamine is a folic acid antagonist it has been associated with bone marrow toxicity while sulfadiazine causes hypersensitivity and allergic reactions up to 20% of population [27-28]. Besides than this conventional treatment has a limited effectiveness, mainly on chronic stage disease, there is not available vaccine for human use.

DHEA is a steroid hormone that is produced, from cholesterol, in the adrenal glands, gonads and brain, and is synthesized from pregnenolone by the action of the 17, 20-desmolase enzyme [29]. It is the most abundant hormone circulating in mammals and can also be a precursor of sexual steroids [30]. The sulphated form of DHEA is majorly found in blood circulation and the free DHEA form (the active form) is only the 3-5 % of the total concentration. Although DHEA is a hormone produced by the organism, it has been postulated for its therapeutic usage as a parasiticide agent. In vitro, low concentrations of DHEA inhibit proliferation, adhesion and motility of *Entamoeba histolytica* trophozoites, while high concentrations induce the lysis of the parasite [31]. DHEA reduces 75% the reproduction of *Taenia crassiceps* cysticercus, in vitro; and in murine model infected with metacestodos of *Taenia* the parasitic charge was 50% reduced when mice were previously treated with the hormone [32]. In a toxoplasmosis acute infection model, DHEA was administrated, pre and pos-infection to immunosupressed mice; DHEA reduced mortality in a 65 % in the pre-treated mice and in a 50 % in post-treated mice; besides this treatment reduced the number of brain cysts in pre-treated infected mice in 90 % and in post-treated mice in 60 % when compared to the control [33]. The effect of estradiol and progesterone has been studied in extracellular tachyzoites of *T. gondii*, showing that estradiol exposure increases the intracellular calcium concentration via the acidic organelles, which increases the secretion of MIC2 and gliding motility in consequence. Progesterone exposure increases the intracellular calcium concentration via the neutral organelles, presenting a contrary effect than observed with estradiol exposure. Although these hormones are able to trigger the calcium signalling in *T. gondii* tachyzoites, none receptor has been reported so far [34].

The development and research of new drugs against toxoplasmosis is relevant; in the search of new therapies with practical application, our research group has studied the effect of sex steroid hormones on the immune response to different parasitic infections. In the present work, we assessed the effect of DHEA, alone or in combination with the conventional treatment S-P, on *Toxoplasma* extracellular tachyzoites. Our results suggest that DHEA could be recognized by a cytochrome b5 family heme/steroid binding domain-containing protein inducing a reduction of passive and active invasion by the modulation of the expression of proteins that are essential during the invasion process, as well as some virulence factors.

## Results

### The treatment with DHEA decreases the viability of *Toxoplasma gondii* extracellular tachyzoites

In order to known the effect of dehydroepiandrosterone hormone on the viability of *Toxoplasma* we exposed extracellular tachyzoites to increasing DHEA concentrations for 30 minutes and two hours. In viability assay, all micromolar concentrations used induced a considerable decrease in the parasite in both times tested. At 30 min, a decrease of between the 25 to 40 % was observed for the 1, 10, 50, 80, 100, 200 and 400 μM concentrations and the maximum effect was observed with 600 μM of DHEA, which reduced the viability in approximately 55% (Fig 1A, grey bars). At two hours, a decrease of the viability of 45 % was observed for 1 and 200 μM concentrations, and the maximum effect was observed for the 10, 100, 400 and 600 μM concentrations that reduced viability in approximately 58 to 62 % (Fig 1A, white bars). These results suggest that the viability of extracellular tachyzoites of *Toxoplasma* is compromised when they are exposed to therapeutic concentrations of DHEA.

**Fig 1.**
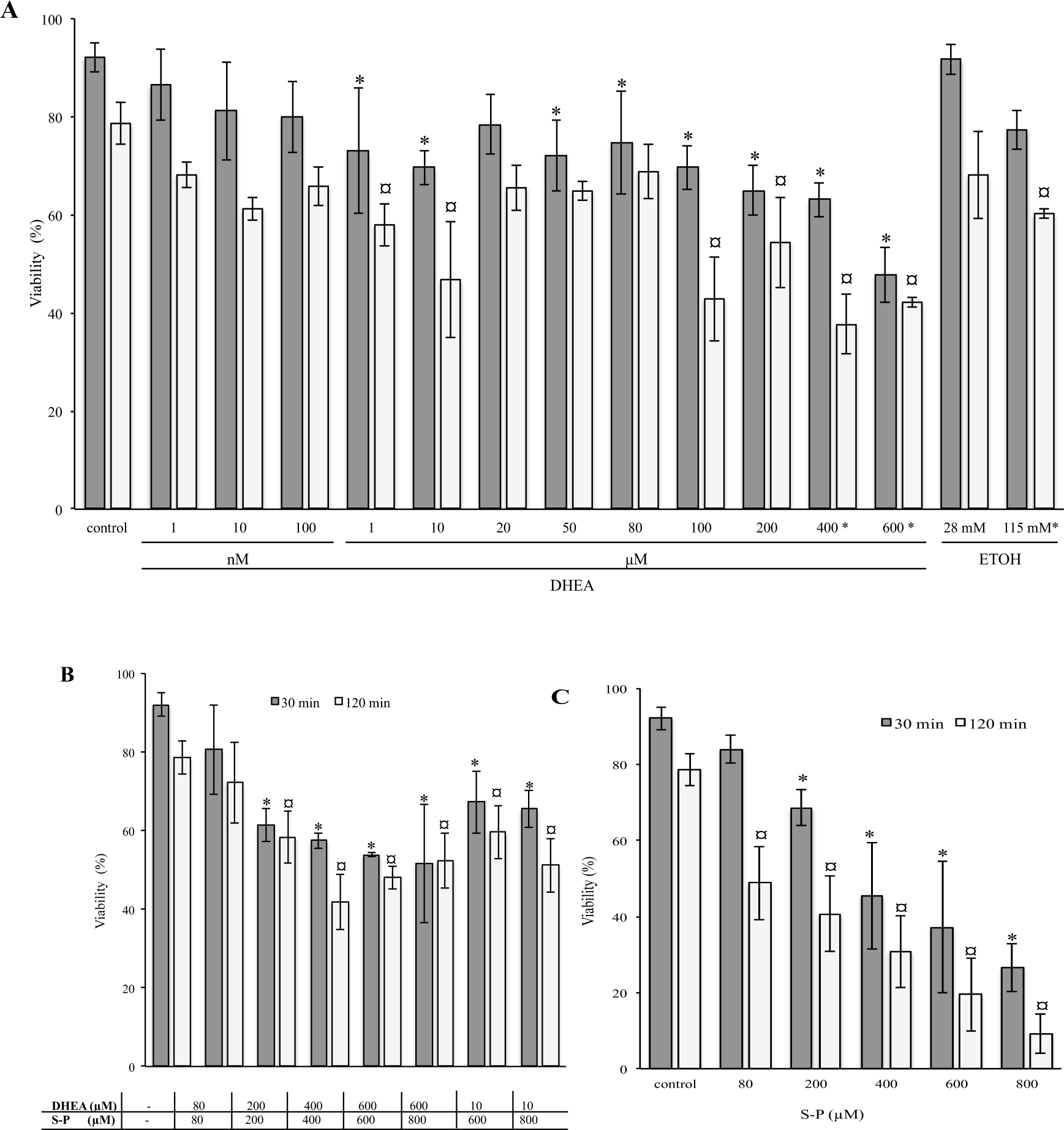
Effect of DHEA on *T. gondii* extracellular tachyzoites viability. A) DHEA B) DHEA/S-P and C) S-P treatment, in X axis showed final concentration of each drug; in Y axis = percentage of viability. Grey bars indicate 30 min and white bars correspond to 2 h of treatment. Control, tachyzoites without treatment in PBS; ETOH correspond to DHEA solution vehicle (ethanol 2.8 and 11.5 μL / 100 μL; * in Y axis, indicate the concentration correspond to high quantity of ethanol used). (*, ¤) Statistical significance compared to the control according to exposure time. P<0.0001

### The combined treatment with DHEA / S-P presents a greater diminution of the viability of *Toxoplasma gondii* extracellular tachyzoites than the individual effect of either treatment

There are the possibility of DHEA could be used as an auxiliary compound in the treatment against Toxoplasma infection, we tested the effect of the conventional treatment with S-P combined with DHEA. First, we used equal concentrations of both drugs at 30 minutes and two hours (Fig 1B). We observed a considerable diminution of the viability with 200 μM concentration in both times tested. At 30 minutes, a decrease between the 40 to the 50 % of the viability was observed since 200, to 600 μM concentrations (Fig 1B, grey bars). The effect observed at 200 μM resulted approximately 10 % higher than the effect observed at the same concentration in the S-P treatment (Figs 1B and 1C, grey bars). In contrast, at two hours the parasite viability was reduced approximately 40 % with the same concentration (Fig 1B, white bars); this reduction are in concordance with reduction induced by DHEA alone; however, the S-P treatment have a better effect inducing a 20 % viability decreased (Figs 1A and 1c, white bars). All the other tested concentrations showed a similar effect (Fig 1B, white bars).

Parasites were also treated with a constant concentration of 10 μM of DHEA combined with 600 or 800 μM of S-P for 30 minutes and two hours. At 30 minutes, both combinations of concentrations showed a reduction of the viability of approximately 35 % (Fig 1B, grey bars); this effect is similar for 10 μM DHEA alone treatment. However, the conventional treatment (S-P) showed a better effect on viability that these modality of combination at 30 min (10 μM DHEA/600 and 800 μM S-P) (Fig 1A vs 1C, grey bars). At two hours, the effect for both combinations of concentrations is lower than individual treatments (Fig 1B vs 1C, white bars).

### The treatment with DHEA and DHEA/S-P reduces the active invasion process

After to treat the tachyzoites with DHEA and DHEA/S-P at several concentrations, we analyse if these parasites were capable to penetrate their human host cell. Hep-2 cells monolayers were exposed at pretreated tachyzoites during 30 min or 2 h (Fig 2). At 30 min, the tachyzoite invasion capacity was inhibit in around 60% respect to the control without treatment, in almost all DHEA concentrations used (asterisks, Fig 2A, grey bars). While, tachyzoites pretreated for 2 h, exhibited a 70% approximately of decrease in invasion process when they were treated with 1, 200 or 400 μM of DHEA, respect to the control without treatment. The maximum effect was observed with 600 μM DHEA, reducing the active invasion in approximately 90 % vs tachyzoites without treatment (Fig 2A, white bars). While DHEA/S-P treatment, only presented significant differences, a decrease of around 50% at 600 μM of both drugs for 30 min (Fig 2B, grey bars); and at 400 μM DHEA/S-P for 2 h, reach approximately a diminution of the active invasion 75 % respect to the parasites without treatment (Fig 2B, white bars). The treatment of the extracellular tachyzoites with S-P had not significant differences in the active invasion independently the concentrations and times tested (Fig 2C).

**Fig 2.**
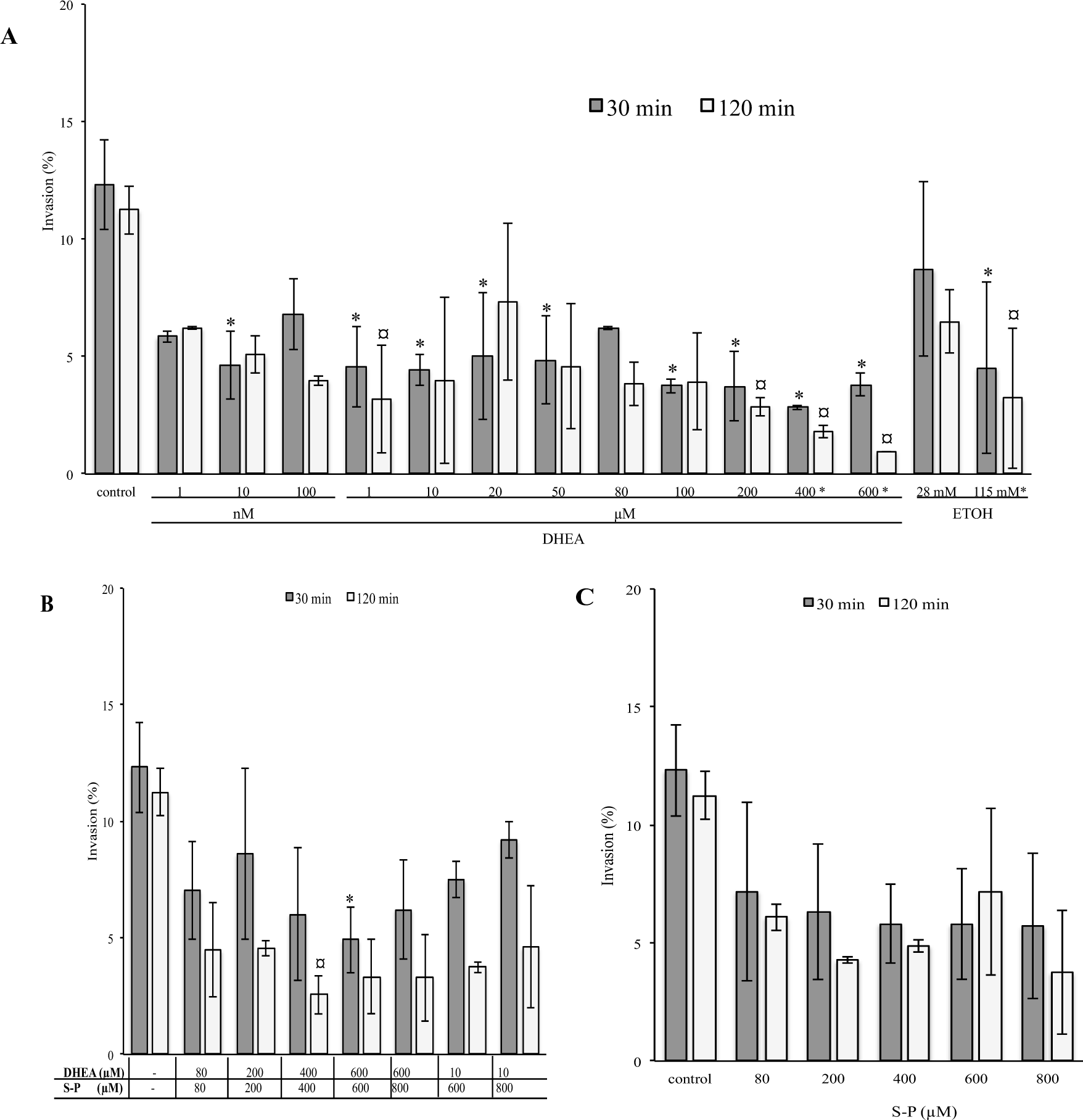
Effect of DHEA in the active invasion process. A) DHEA B) DHEA/S-P and C) S-P treatment, X axis = final concentration of each drug; Y axis = percentage of HEp-2 cells that contained at least one parasitophorous vacuole in the cellular cytoplasm. Grey bars, 30 min and white bars, 2 h of treatment; control, PBS ETOH correspond to DHEA solution vehicle (ethanol 2.8 and 11.5 μL / 100 μL; * in Y axis, indicate the concentration correspond to high quantity of ethanol used). (*, ¤) Statistical significance compared to the control according to exposure time. P<0.0001

### The treatment with DHEA reduces the passive invasion process

*T. gondii* have the capacity to invaded all nucleated cells, to introduce at phagocytic cells like macrophages, it used the active machinery of the phagocytic cells themselves, this kind of invasion is called passive. In order to know if DHEA has an effect in this process we put together fresh macrophages with pretreated extracellular tachyzoites for 30 min or 2 h (Fig 3). First, we compared the percentage of phagocytized untreated tachyzoites by LPS-activated macrophages against the percentage of phagocytized untreated tachyzoites by non-activated macrophages. Activated macrophages phagocytized approximately 40% more untreated tachyzoites compared with non-activated macrophages (Fig 3A, activated vs non-activated). For this reason we use activated macrophages for the consecutive assays. Passive invasion was reduced in approximately 35 % when activated macrophages were exposed to pretreated tachyzoites with 600 μM DHEA for 30 min (Fig 3A, grey bars), and to pretreated tachyzoites with 80, 400 and 600 μM DHEA for 2 h (Fig 3A, white bars). The combined (DHEA/S-P) and the conventional (S-P) treatments on extracellular tachyzoites have no effect in the passive invasion, independently of the concentrations and times (Fig 3B and 3C, respectively).

**Fig 3.**
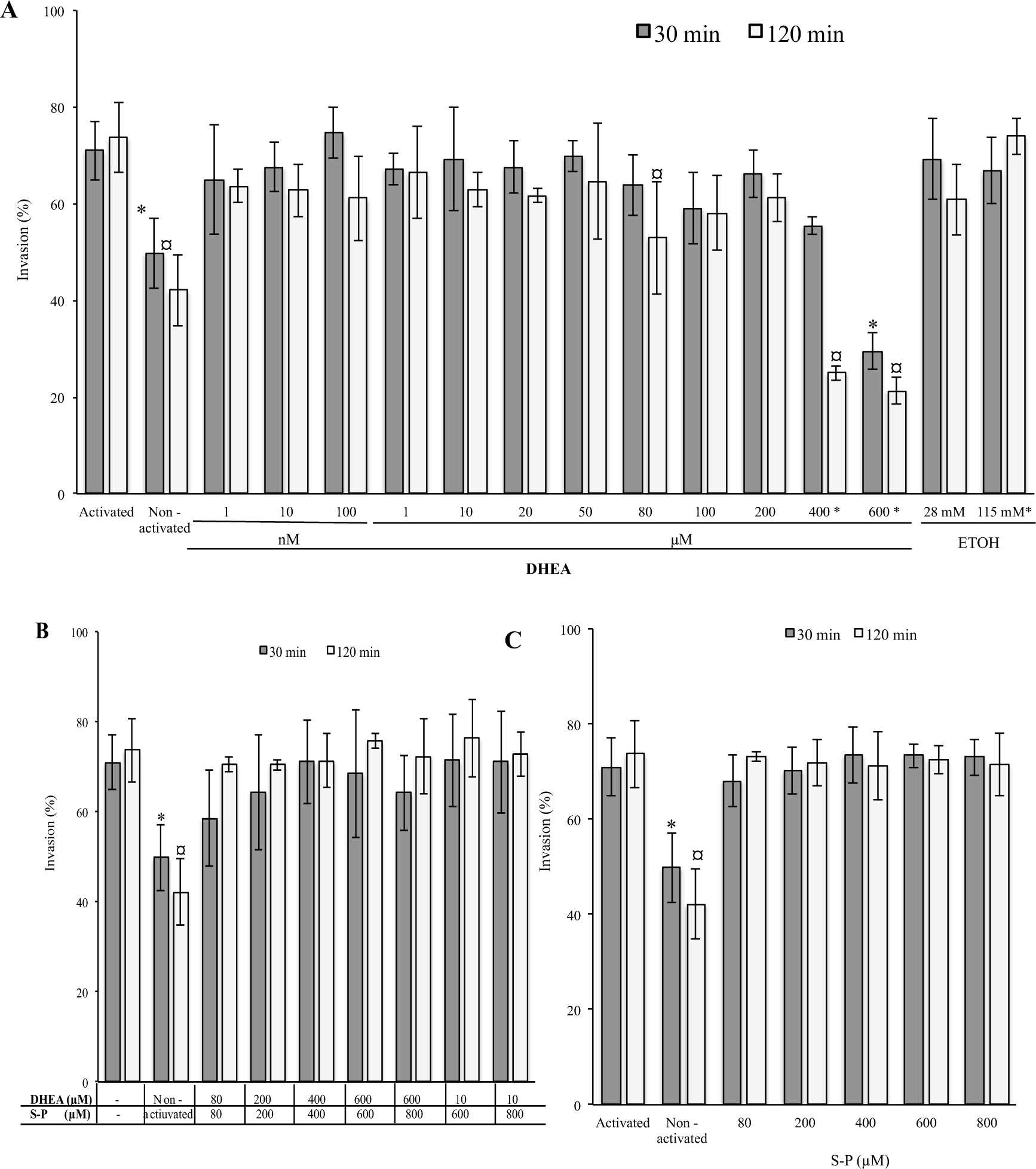
Effect of DHEA in the passive invasion process. A) DHEA B) DHEA/S-P and C) S-P treatment, X axis = final concentration of each drug; Y axis = percentage of macrophages that contained at least one PV in the cellular cytoplasm. Grey bars, 30 min and white bars, 2 h of treatment; control, PBS ETOH correspond to DHEA solution vehicle (ethanol 2.8 and 11.5 μL / 100 μL; * in Y axis, indicate the concentration correspond to high quantity of ethanol used). (*, ¤) Statistical significance compared to the control according to exposure time. P<0.0001

### Tachyzoites treated with DHEA and DHEA / S-P present morphological changes

We analysed if the decrement in invasion process could be related to morphological changes induced by the DHEA treatment on extracellular tachyzoites. The ultrastructure images of extracellular parasites treated as in the viability assay, for all concentrations of each treatment, DHEA and S-P alone and DHEA/S-P, were obtained by TEM (Figs 4 and 5). The untreated and vehicle control (ethanol) tachyzoites, are showed in Fig 4A and 4B, respectively. The DHEA treatment at 10 μM – 30 min, preserves all the typical structures such as micronemes, rhoptries, dense granules, nuclei and mitochondria.

**Fig 4.**
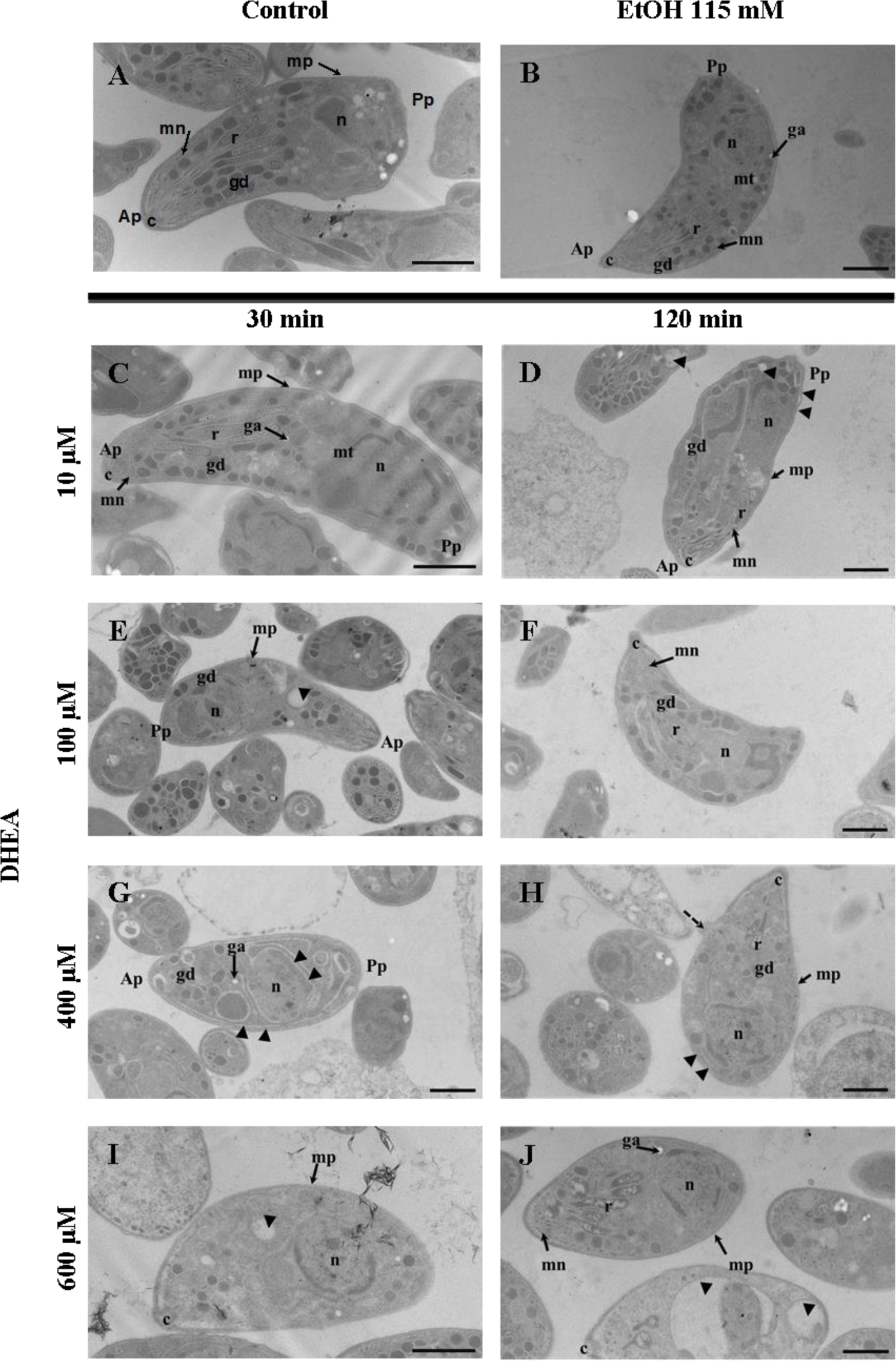
Effect of DHEA in the ultrastructure of *Toxoplasma gondii* extracellular tachyzoites. A) Morphology of extracellular tachyzoites untreated and B) tachyzoites exposure to ethanol, corresponding at 2h of exposure. Extracellular tachyzoites exposure to C) 10 μM, E) 100 μM, G) 400 μM and I) 600 μM final concentration of DHEA for 30 min; or D) 10 μM, F) 100 μM, H) 400 μM and J) 600 μM final concentration of DHEA for 2 h.. c, conoid; r, rhoptries; mn, micronemes; gd, dense granules; ga, amylopectin granules; n, nuclei; m, plasma membrane; re, endoplasmic reticulum; mt, microtubules; Ap, apical pole; Pp, posterior pole Bars= 1 µm.

**Fig 5.**
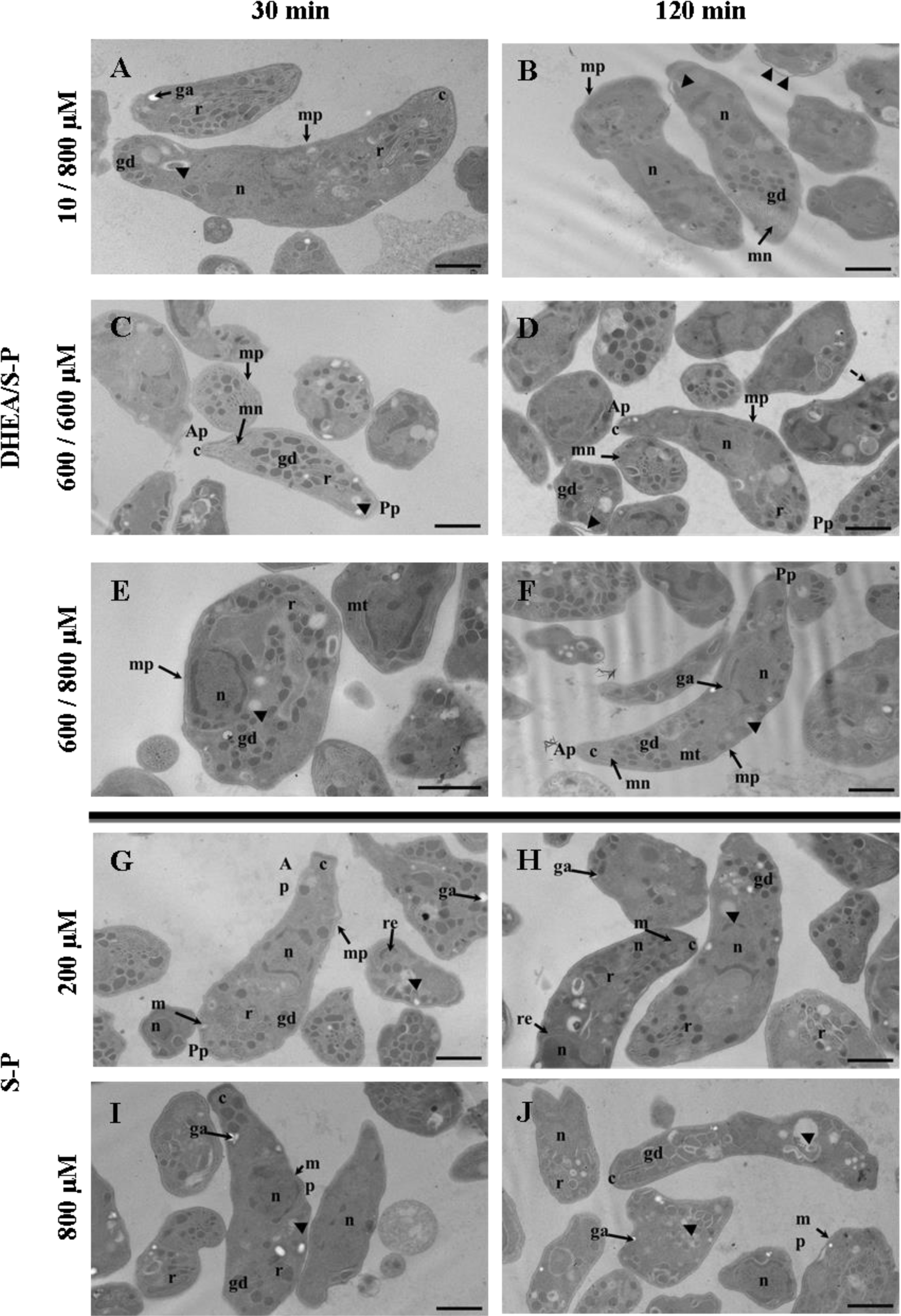
Effect of DHEA / S-P in the ultrastructure of *Toxoplasma gondii* extracellular tachyzoites. Extracellular tachyzoites treated with A) 10 / 800 μM, B) 600 / 600 μM, C) 600 / 800 μM final concentration of DHEA / S-P, respectively for 30 minutes or D) 10 / 800 μM, E) 600 / 600 μM, F) 600 / 800 μM final concentration of DHEA / S-P, respectively for 120 minutes. c, conoid; r, rhoptries; mn, micronemes; gd, dense granules; ga, amylopectin granules; n, nuclei; m, plasma membrane; mt, microtubules; Ap, apical pole; Pp, posterior pole. Bar = 1 µm.

Interestingly, we were capable to observe the presence of granules apparently of amylopectin, which are exclusive of bradyzoite stage; and some areas of plasmatic membrane look wavy (Fig 4C). Importantly, the effect of DHEA on tachyzoites structure is related to the concentration and time used. At 100 μM DHEA for 30 min, the parasites seem to lose their typical half-moon shape and some of them present amylopectin granules (Fig 4E). While with 400 μM DHEA (30 min), parasites look a little swollen and present bigger amylopectin granules (Fig 4G). At 30 minutes with 600 μM DHEA, some parasites seem to preserve their half-moon shape, with a reorganization of the organelles and the presence of amylopectin granules; however, at this concentration of DHEA there are a greater amount of phantom structures in the samples (Fig 4I). Longer time of DHEA exposition induced greater changes in the extracellular tachyzoites morphology. Parasites exposure to 10 and 100 μM of DHEA by two hours, showed an amoeboid shape with a totally lost of the intracellular organization and the apical polarity; the dense granules lost their circle shape and the presence of some amylopectin granules were observe to (Fig 4D and 4F). In the same time but exposure with 400 and 600 μM of DHEA, parasites look inflated with a balloon shape, although they seem to preserve organelles typical organization, as well as the apical polarity (Fig 4H and 4J).

In the combined treatment, DHEA/S-P at 10/800 μM, 600/600 μM and 600/800 μM, the effect was observed since 30 min of exposure and it was consistent after 2 h (Fig 5). At 10/800 μM DHEA/S-P the parasites lose their typical shape and present amoeboid and elongated shape, they also start to lose the intracellular organization and present amylopectin granules (Fig 5A-B). At higher concentrations of DHEA/S-P 600/600 μM, some parasites preserved the typical shape lose the organization of the organelles, finding the rhoptries at the posterior pole, the dense granules lost their circled shape, and they present amylopectin granules (Fig 5C-D). Finally, with 800/800 μM of combined treatment, the appearance of tachyzoites was amorphous with loss of the organelles organization and again, presence of amylopectin granules (Fig 5E-F)

The conventional treatment induced important changes in the extracellular tachyzoites, independently of concentration or time exposure. At 200 and 800 μM of S-P for 30 min and 2 h, the tachyzoites loose their typical shape and was possible find tachyzoites with amoeboid, elongated or amorphous shape, they present amylopectin granules, loose the apical polarity and the organization of the organelles (Fig 5G-J).

### Treatment with DHEA, S-P or DHEA / S-P induces changes in the protein expression since 30 minutes of exposure

To determine the effects that the treatments with DHEA, S-P and DHEA / S-P have at molecular level, we used in extracellular tachyzoites to compare the protein profile of the treated parasites against the protein profile of tachyzoites without treatment (Fig 6). The protein profile of the control exhibited 159 spots (Fig 6A); the group treated with DHEA 10 μM for 30 min exhibited 165 spots, which 105 were match with control (Fig 6B); the group treated with S-P 800 μM for 30 min exhibited 126 spots, which 99 were match with control (Fig 6C); while the group treated with DHEA / S-P 10 / 800 μM for 30 min exhibited 213 spots, which 113 were match with control (Fig 6D). Protein profiles were analyzed as described in methods and materials section, and we selected the proteins that showed greater changes in their expression between the treatments and respect to the control. Thirty proteins were identified by its molecular weight and isoelectric point (Table 1), as described in methods and materials section, and were classified by their probable location (Fig 7A). We observed that most of the proteins that change their expression are dense granules proteins, followed by proteins from plasma membrane and cytoplasm (Fig 7A). Then, we graphed the proteins that diminishes their expression respect to the control (Fig 7B), and the proteins that maintain or increase their expression respect to the control (Fig 7C). We can observe that treatment with DHEA leads in a diminution of the expression of proteins from dense granules, micronemes, apicoplast, peroxisome, plasma membrane, mitochondria and cytoskeleton; while proteins from cytoplasm increases their expression (Fig 7 insets B and C, light grey bars). S-P treatment provokes the reduction in the expression of proteins from dense granules, micronemes, cytoplasm, apicoplast, peroxisome, plasma membrane and mitochondria; and the increment of proteins from rhoptries, IMC and cytoskeleton (Fig 7 insets B and C, dark grey bars). Combined treatment with DHEA / S-P, induces a lower expression of proteins from dense granules, micronemes, cytoplasm, apicoplast and peroxisome; and a higher expression of proteins from rhoptries, plasma membrane and mitochondria (Fig 7 insets B and C, black bars).

**Table 1.**
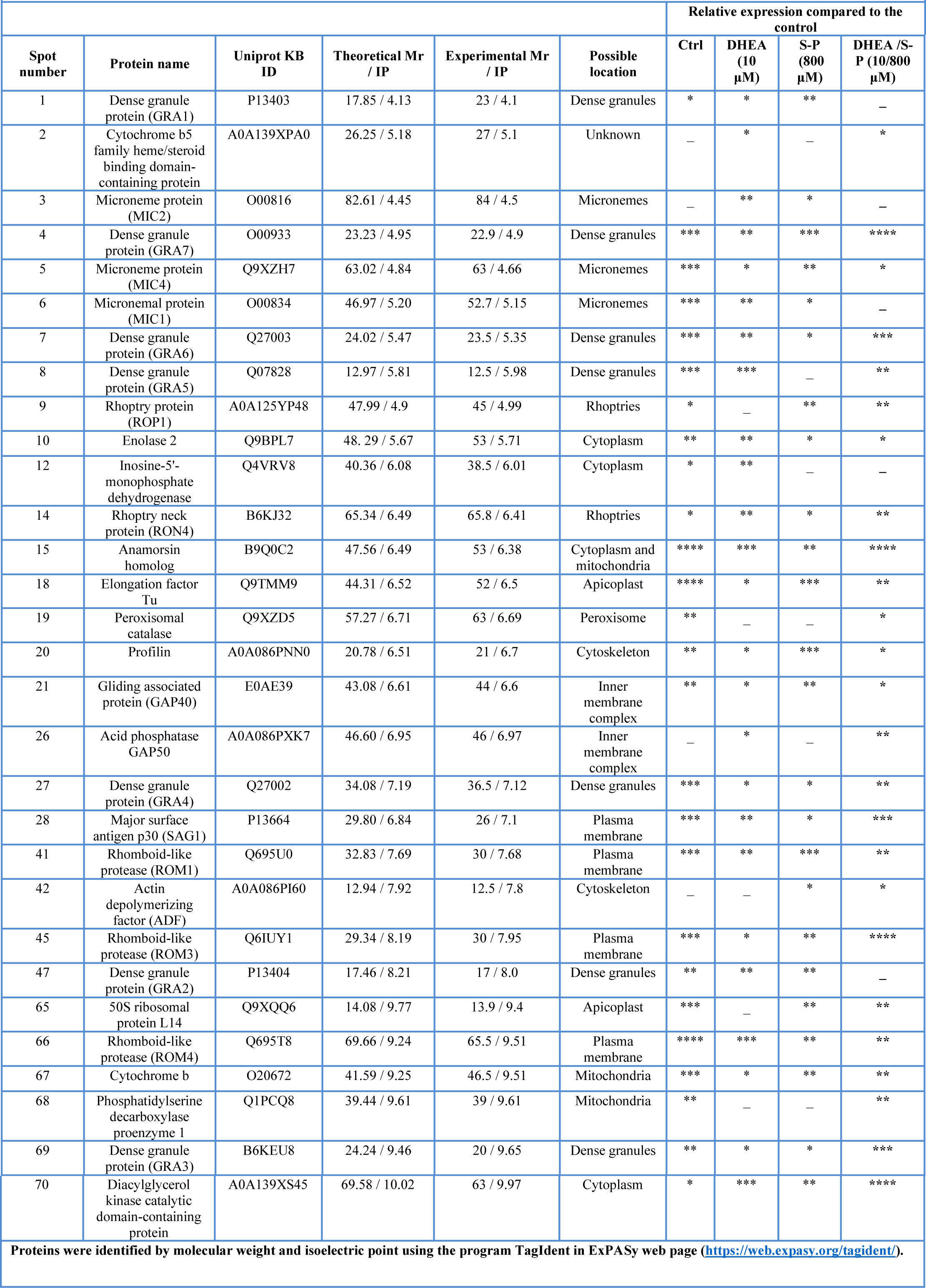
Protein differential expression in whole extract of extracellular tachyzoites.

**Fig 6.**
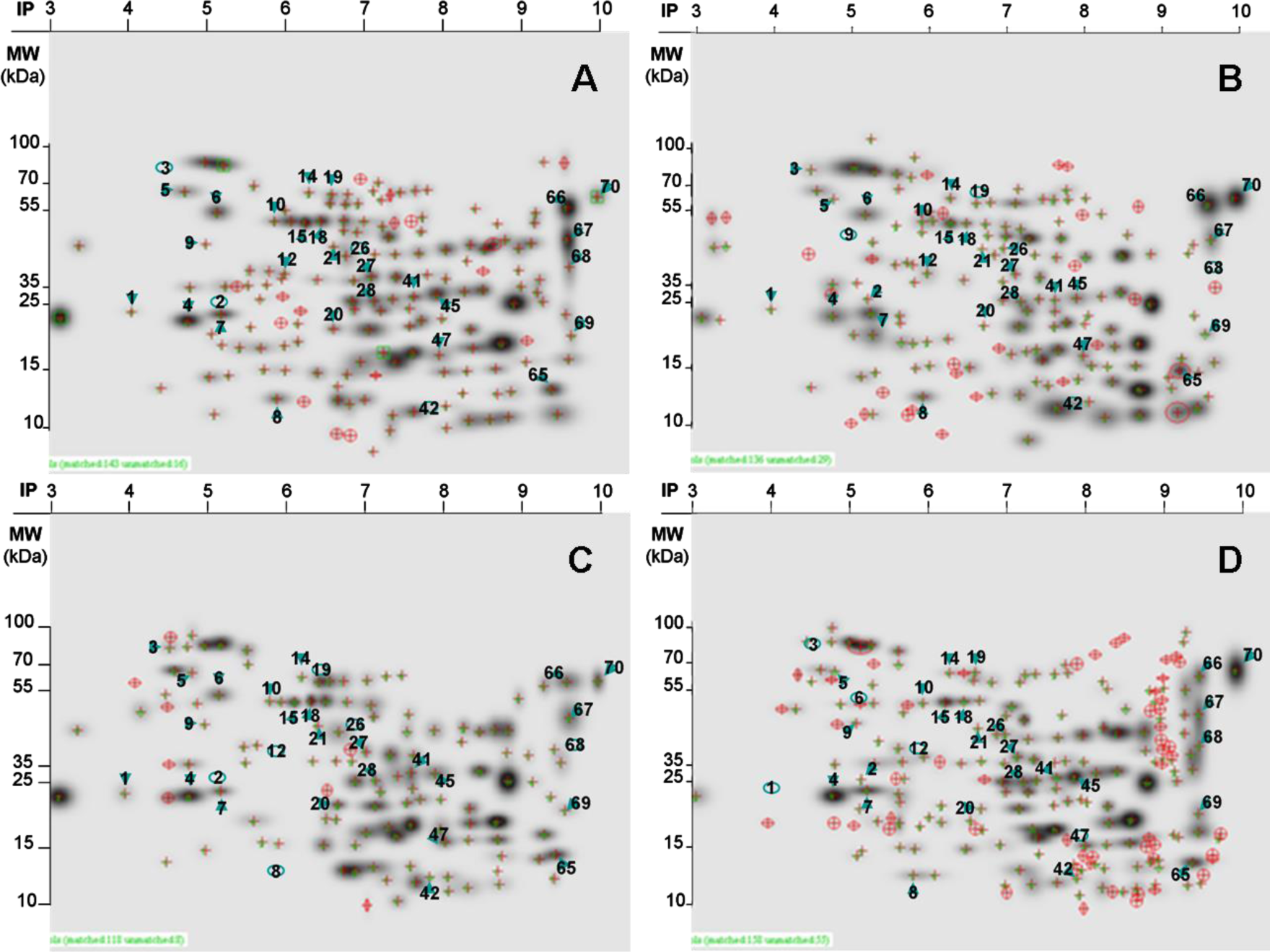
Proteomic profile of whole extract of extracellular tachyzoites. 30 minutes of treatment with: A) Control without treatment; B) DHEA 10 μM; C) S-P 800 μM and D) DHEA / S-P 10 / 800 μM. Arrowheads point out the spots (identified in table 1) that were identified in all treatments; the spots that are absent in treatments respect to the control are circled.

**Fig 7.**
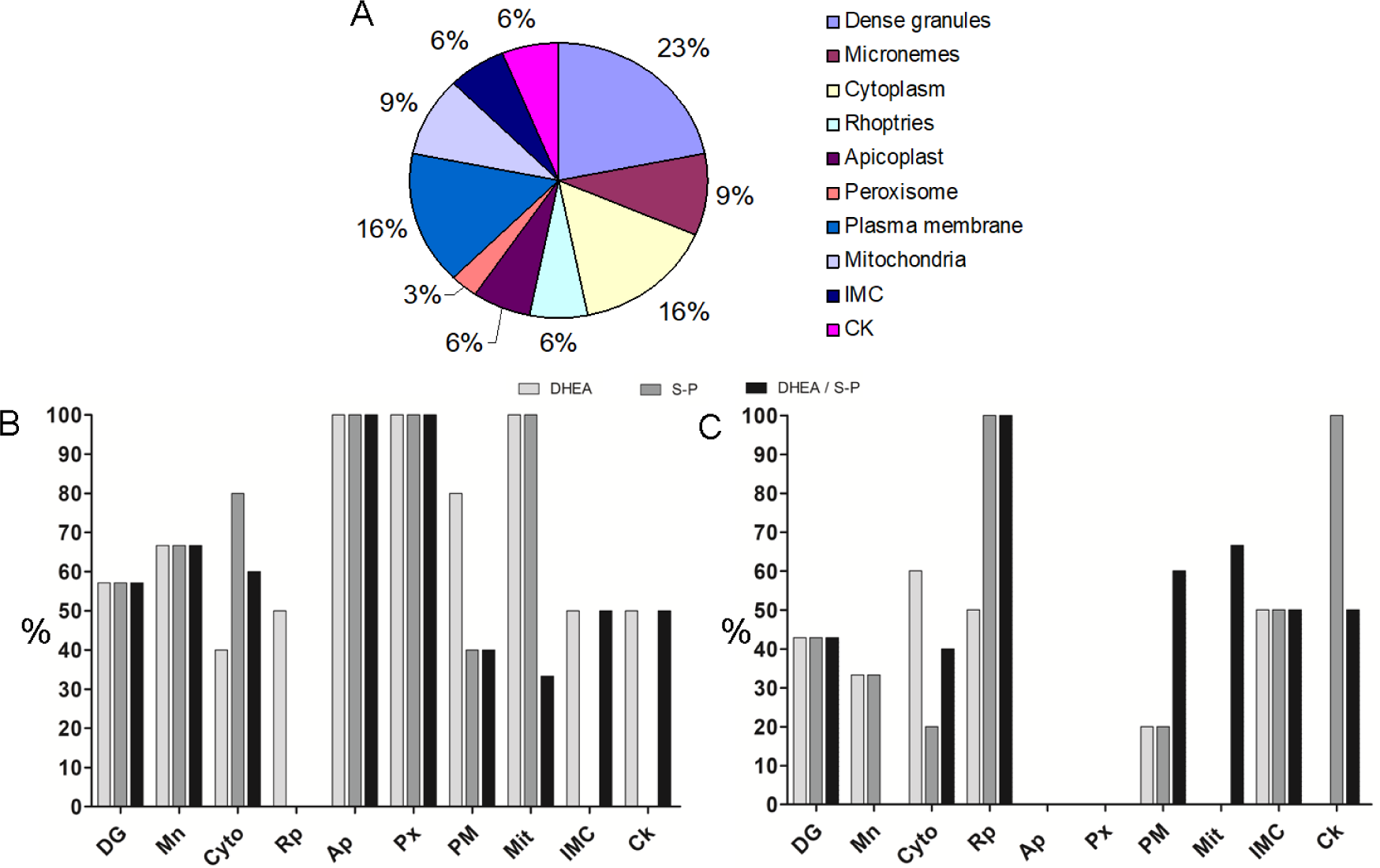
Classification of proteins that exhibited differential expression. A) Proteins that change their expression after 30 minutes of treatment with DHEA 10 μM, S-P 800 μM or DHEA / S-P 10 / 800 μM, grouped by their probable location. B) Percentage of proteins that decrease their expression respect to the control without treatment (y axis). C) Percentage of proteins that maintain or increase their expression respect to the control without treatment (y axis). X axis, probable location; DG, dense granules; Mn, micronemes; Cyto, cytoplasm; Rp, rhoptries; Ap, apicoplast; Px, peroxisome; PM, plasma membrane; Mit, mitochondria; IMC, inner membrane complex; Ck, cytoskeleton.

### Interaction of DHEA with the Cytochrome b5 family heme/steroid binding domain-containing protein

Interestingly the spot number 2, which exhibited an experimental molecular weight of 27 KDa and isoelectric point of 5.1, and that is expressed only in the protein profile of tachyzoites treated with DHEA; was identified as a cytochrome b5 family heme/steroid binding domain-containing protein with a theoretical molecular weight of 26.25 KDa and isoelectric point of 5.18. The primary sequence of the protein was aligned in the NCBI web site with the BLAST program (https://blast.ncbi.nlm.nih.gov/Blast.cgi?PROGRAM=blastp&PAGE_TYPE=BlastSearch&LINK_LOC=blasthome) and the protein was aligned with other steroid binding proteins from other Eukaryotes (S1 figure A-C), then we aligned only the domain section (aminoacids from 129 to 176) and the result was similar (S1 figure D-F).

The *T. gondii* model generated is the best that can be obtained given that the only available template has a 36.9% homology; this corresponds to the 111 residues located at the carboxyl-terminus of the full 243 residue protein. Most of the residues know to interact with the heme group in the PGRMC1 structure are identical in our predicted structure for *T. gondii* PGRMC. The resulting model has a heme group partially buried and contributing significantly to the binding of all of the ligands tested (Fig 8). In every case, the three best results for each ligand were in contact with the heme group on the surface of the protein. Notably, progesterone is the most tightly bound ligand followed by dehydroepiandrosterone, testosterone and 4-5 alpha dihydrotestosterone (Table 2).

**Table 2.** Ligands that present best affinities to TgPGRMC.

| Table 2. Ligands that present best affinities to TgPGRMC |  |
| --- | --- |
| Ligand | Vina<br>kcal/mol |
| Octanoate | -4.1 |
| Decanoate | -4.4 |
| Dodecanoate | -4.6 |
| Myristate | -5 |
| Palmitate | -4.6 |
| Stearic | -5.1 |
| Oleate | -5.4 |
| Linoleate | -5.5 |
| **DHEA** | **-7.4** |
| **4-5 alpha-Dihydrotestosterone** | **-7.4** |
| Aldosterone | -7.1 |
| Estriol | -7.2 |
| Cholesterol | -6.6 |
| **Progesterone** | **-7.6** |
| **Testosterone** | **-7.4** |
| Corticosterone | -6.8 |
| Beta-estradiol | -6.7 |
| Cortisol | -6.5 |
| Pyrimethamine | -5.9 |
| Sulfadiazine | -5.5 |

**Fig 8.**
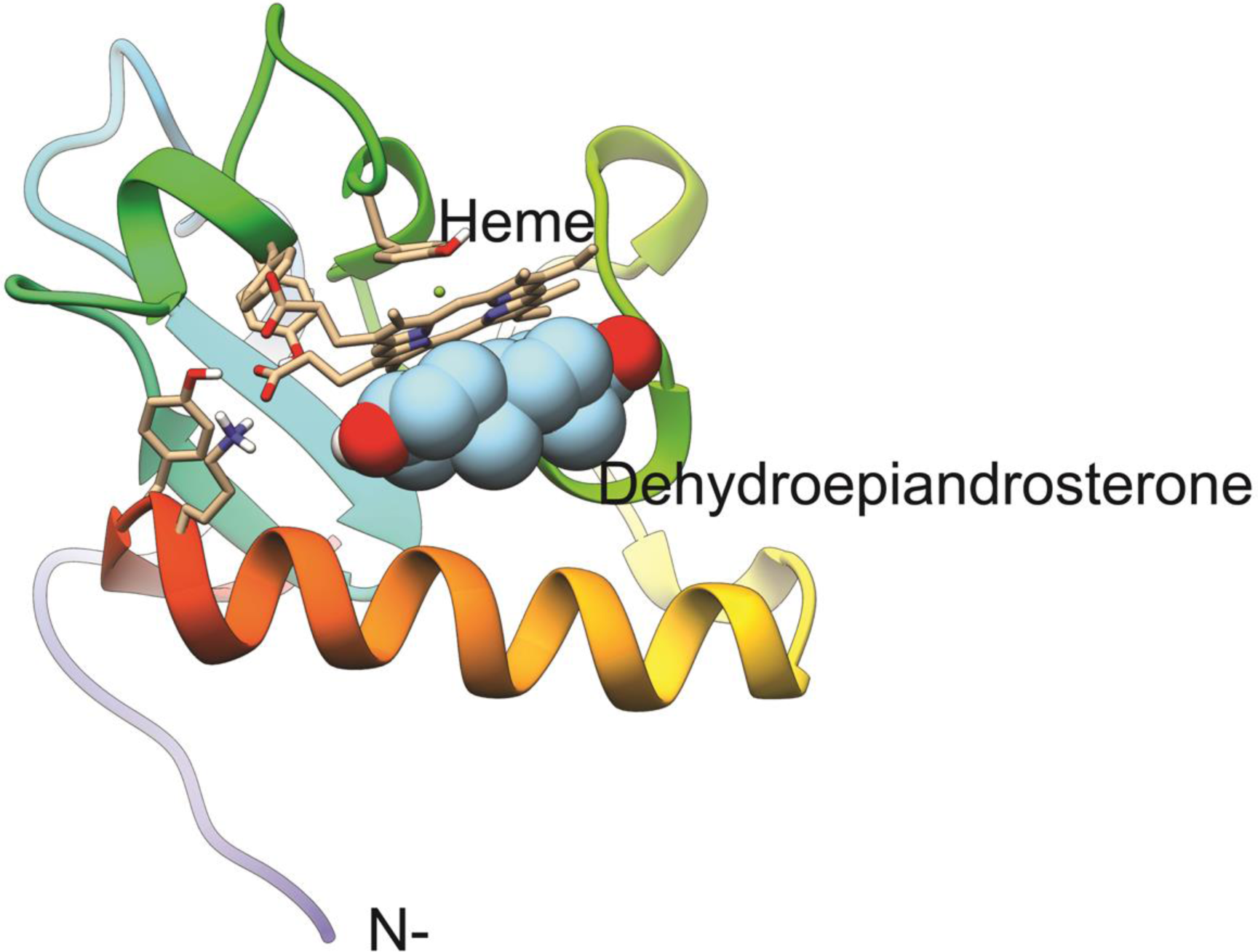
Model for *T. gondii* PGRMC homolog and its docking to DHEA. The model for PGRMC contains a binding pocket for a heme group that functions as the binding site for DHEA. TYR158 binds the heme group on one face while the other binds DHEA, blocking any interaction at that site.

Given that residue TYR158 (numbering based on the whole sequence cloned) provides the fifth coordination to the heme-iron, forcing ligand interaction to occur on the unoccupied side of the heme group.

Pyrimethamine and sulfadiazine were also found to bind the heme group but with significantly lower affinity (bottom of Table 2). Given that these affinities are about 1.5 kcal/mol lower, it is likely that they are non-specific as well as is the case for fatty acids included in the docking.

It is unknown if *T. gondii* PGRMC is able to dimerize as its template (*Homo sapiens* PGRMC1), but the interactions with the ligands tested in the present work would block or compete a similar interaction.

## Discussion

DHEA induces a decrement in the viability of extracellular tachyzoites at 100 μM at both times tested (30 min and 2 h); even if effect was lower than the observed with standard therapy (S-P); results are similar to the obtained when trypomastigotes of *T. cruzi*, another intracellular parasite were treated with 128 μM final concentration of DHEA combined with melatonin for 24 h [35]. Different to the standard treatment, whose response was directly proportional to the concentration, the effect obtained with DHEA exposure was independent on the concentration. This suggests that both drugs have different targets inside of the tachyzoite. In order to evaluate an accumulative effect of the DHEA with S-P we tested the combined treatment, resulting in a similar effect as the obtained when DHEA alone was administered. It is important to mention that previously reported DHEA parasiticide effect depends on the administration scheme, parasite lineage and experimental conditions.

Once we determined that viability was affected by DHEA treatment, we assessed if the hormone has an effect in the invasion process, which is the most important biological process for the establishment and maintenance of the infection; as tachyzoites are able to infect phagocytic and non-phagocytic cells in two different mechanisms that has been defined as passive and active invasion, respectively, we determined the effect of the DHEA in both.

Tachyzoites reduced its ability to invade HEp-2 monolayers, when they were treated with DHEA; this reduction was higher than the observed with the conventional treatment with S-P. Differently to passive invasion, active invasion was reduced when parasites were treated with the DHEA / S-P combination at high concentrations, 400/400 μM at 30 minutes and 600/600 μM at 2 h. During the active invasion, *Toxoplasma* is the effector cell and the recognition of an unknown component in the plasma membrane of the host cell, is required; this event is determined by the GPI anchored proteins from *Toxoplasma*, being the SAG1 protein the most abundant in the plasma membrane of tachyzoites [13]. The expression of SAG1was reduced in parasites that were treated with DHEA and S-P, while the parasites that were treated with DHEA / S-P maintain the expression of this protein similar to the control; is well known that SAG1 is not the only protein that act in this first process of attachment, due to the Sag1^-^ mutants are still able to invade [13]. The motility is essential for the invasion and it depends of the glideosome complex; in this respect, several proteins that participate in the formation of the glideosome and in its correct function reduced its expression when parasites were treated with DHEA and DHEA / S-P, such as GAP40, profilin and ROM4. GAP40 protein act as an anchor for the rest of the glideosome complex [36]; the role of profilin is to sequester the G actin in order to enhance the polymerization, and it has been demonstrated to be essential for gliding motility and cell cycle in *Toxoplasma* [37]; and ROM4 is a rhomboid protease necessary for the cleavage of the complex MIC2 / AMA1 that is formed for the establishing of the MJ during the invasion process, the correct cleavage of the complex is required for correct reorientation of the parasite and gliding motility [38]. The reduction of the expression of these proteins could explain the diminution in the parasite ability for invade the Hep2 cells monolayers when are treated with DHEA or DHEA / S-P.

The analysis in primary cultures of macrophages revealed that high concentrations of DHEA affect the tachyzoites ability to establish in the cytoplasm of the phagocytic cell. In comparison with DHEA treatment, neither the conventional treatment with S-P, nor the combined treatment DHEA / S-P present significant differences, even when combined treatment reduced the extracellular viability. It has been proposed that tachyzoites can transform the phagocytic vacuole into a parasitoforous vacuole by two different processes; the first includes the formation of the moving junction at the same time that the parasite is phagocytated, and the second proposal is that once the parasite has been phagocytated, is able to invade the phagolysosomal vacuole [13, 39]. Both proposals imply the fusion of the tachyzoite plasma membrane with the macrophage plasma membrane or with the phagolysosomal membrane; this mechanism involves the secretion of proteins from secretory organelles, such as MIC2 and RON4 [18]. In an interesting way, the protein MIC2 exhibited a higher expression when extracellular tachyzoites were exposure to DHEA, its expression was lower with the exposure to S-P and undetectable in both, the combined treatment and control; and the expression of RON4 was higher in the both treatments that include DHEA compared to the S-P treatment and the control (Table 1). It has been previously reported that extracellular tachyzoites that were exposure to progesterone, inhibit the secretion of MIC2 but didn’t affect its expression [34]; this inhibition of the secretion leads on the inhibition of gliding motility. In this work we didn’t collected the secretion products, but we must collect in future assays in order to determine if these proteins are secreted or not, nevertheless the increment of MIC2 could due to a failure on its secretion.

Complementary assays (not shown) revealed that even if the parasites were established inside of the macrophages, the evasion of the lysis was inhibited; this suggest that DHEA treatment could prevent the block of the phosphorylation of the host Immune-Related GTPases (IRGs) by ROP18 and GRA7 proteins from the parasite, diminishing its ability to escape of lysosomal degradation. Concordantly with this, the expression of GRA7 was reduced when parasites were treated with DHEA, while S-P treatment exhibited a similar expression to the control and in an unexpected way the combined treatment with DHEA / S-P increased the expression of the protein (Table 1). GRA7 interacts with the ROP18 kinase in a complex that targets the host IRGs mediating macrophage survival and acute virulence; ΔGRA7 strain reduces the virulence in a half and the parasites can’t evade the lysosomal degradation [40].

The effect of the DHEA in the structure of the extracellular tachyzoites resulted in the alteration of the plasmatic membrane structure, the organization of the organelles, the structure of the secretory organelles and cytoskeleton structures. At the higher concentrations (400 and 600 μM) tested for 2 h, tachyzoites looked totally swollen; tachyzoites that were treated with S-P and DHEA / S-P, shown the same structure alterations, at lowest concentration and time tested, except for the swollen shape. The lost of the structure and location of secretory organelles when parasites were treated with DHEA, is consistently with the reduction in the invasion and in the ability to escape of the macrophage lysis; because both mechanisms depend on the secreted proteins from micronemes, rhoptries and dense granules; this effect is also related with the diminution in the expression of these proteins, as was previously discussed. Additionally, GRA3 expression was reduced when parasites were exposed to DHEA and S-P; recently was reported that GRA3 could has a role in the stabilization of the subpellicular cytoskeleton network, as ΔGRA3 strain tachyzoites purified cytoskeletons lost the organization of this structure [41].

Another three proteins with differential expression regulation worth noting are of diacylglycerol kinase catalytic domain-containing protein, enolase 2 and a cytochrome b5 family heme/steroid binding domain-containing protein. The former, is a protein that is essential for the micronemes correct secretion [42]; this protein increases its expression in all three treatment schemes, as we didn’t collect secretion products more experiments should be achieved in order to determine the effect of the hormone in the function of this protein.

Enolase 2 besides of being specific of the tachyzoite stage, acts as a transcription factor during intracellular proliferation [43-44]; this protein maintains its expression similar to the control when parasites were expose to DHEA, while its expression was reduced with the S-P and DHEA / S-P treatment. We didn’t identify proteins specific of bradyzoite stage, but would be interesting the search of some proteins in order to determine a possible transformation from tachyzoite to bradyzoite stage.

The later protein was expressed only in the parasites that were exposed to DHEA. Given its homology to PGRMC1 and 2, proteins known for their roles as progesterone receptors as well as interactions with the family of cytochromes P450 monooxygenase systems (doi.org/10.3389/fphar.2017.00159), it is not surprising to find it associated to a drug metabolism and response function in *T. gondii*. Interaction between this PGRMC homolog and DHEA could potentially block normal activating interactions with CYPs thus preventing the removal of the steroid. Thus, both the experiment in cells, as well as the molecular docking provides evidence for dehydroepiandrosterone to have a different target and effect than S-P.

Together our results suggest that DHEA can be proposed as a new treatment by itself or in a combined scheme with conventional treatments; however more experiments should be achieved in order to investigate its parasiticide effect *in vivo*.

## Conclusion

DHEA parasiticide effect could be due to its interaction with the cytochrome b5 family heme/steroid binding domain-containing protein. DHEA treatment reduces the expression of proteins that are essential for the motility and virulence of RH strain tachyzoites and it is likely to block removal of DHEA by CYPs. This leads on an alteration of the ultrastructure of the parasites, the lost of the organelles organization, the lost of structure of the secretory organelles and of the cell shape. These alterations induce a reduction of the invasion ability of the tachyzoites to HEp-2 monolayers; a reduction of the ability to scape of the macrophage’s lysis; and a reduction of the viability *in vitro*.

## Materials and methods

### Drugs, reagents and solutions

The DHEA (Sigma chemical Co. Steinheim, Germany) was dissolved in anhydrous ethanol (Sigma chemical Co. Steinheim, Germany). The sulfadiazine-pyrimethamine was obtained in its commercial formulation (Bactropin ®, trimethoprim-sulfamethoxazole 160/800 mg).

### Animals

Male Balb-C/Ann mice, between 6-8 weeks of age, used for parasite infection were maintained in a pathogen-free environment with regulates conditions of temperature, humidity and filtered air. Animals were fed with autoclaved food and water at libitum; and maintained according to the Mexican Federal Regulations for Animal Production, Care and Experimentation (NOM-062-ZOO-1999, Ministry of Agriculture; Mexico City, Mexico). All efforts to minimize animal suffering and to reduce the number of animals used were made.

### Maintenance and purification of *T. gondii* tachyzoites

Tachyzoites of RH strain was maintained by intraperitoneal (ip) passages in Balb/cAnN mice. After cervical dislocation parasites were recovered from peritoneal exudates, washed with 1X PBS (138 mM NaCl, 1.1 mM K2PO4, 0.1 mM Na2HPO4 and 2.7 mM KCl, pH 7.2) and purified by filtration through 5 μm pore polycarbonate membranes (Merck Millipore Co. Cork, Ireland).

### HEp-2 cell culture

Hep-2 cells (CCL-23), derived from Human larynx epidermoid carcinoma epithelial cells were obtained from American Type Culture Collection (ATCC). The cellular culture was maintained in Minimal Essential Medium media (MEM, Gibco, Thermo Fisher, NY, USA), supplemented with 8 % fetal bovine serum (FBS Gibco, Thermo Fisher, NY, USA) and 1 % PES, under 5 % CO2 atmosphere at 37 °.

### Murine macrophages culture

Sterile mineral oil (1 mL) was inoculated in the peritoneum of male BalbC/AnN mice, after five days the mice were sacrificed and intraperitoneal macrophages were recovered using 1% glucose-PBS solution; macrophages were centrifuged and pellet was resuspended in Dulbecco’s Modified Eagle Medium (DMEM, Gibco, Thermo Fisher, NY, USA), supplemented with 8% of fetal bovine serum (FBS) and 1% of penicillin-streptomycin (PES 10, 000 u/mL, Thermo Fisher, NY, USA). Macrophages were seeded over sterile coverslips in a ratio of 250 × 103 / cm2 and they were maintained at 5% CO2 atmosphere at 37 °.

### Macrophage activation

Macrophages were washed with fresh DMEM 24 h after they were seeded. Lipopolysaccharides (LPS, Sigma chemical Co. Steinheim, Germany) were added at 30 ng/mL final concentration for 1 h, in order to activate the macrophages. Then they were invaded with tachyzoites as described in “Invasion assays” section.

### Viability of extracellular tachyzoites

Purified parasites (6 × 106 cells) were exposure to increasing final concentrations of DHEA (1, 10 and 100 nM, and 1, 10, 20, 50, 80, 100, 200, 400 and 600 μM), of Sulfadiazine-Pirimetamine (80, 200, 400, 600 and 800 μM) and to combined treatment DHEA/ S-P (80/80, 200/200, 400/400, 600/600, 600/800, 10/600 and 10/800 μM), all drugs were diluted in PBS to final concentrations. The exposition was held for 0.5 or 2h at room temperature (RT) with gentle agitation. The ratio between live and death tachyzoites was measured by exclusion technique with trypan blue (4% Gibco, Thermo Fisher, NY, USA), 300 parasites were counted under an optical microscope (AxioObserve Microscope, Carl Zeiss Mexico). This assay was realized by triplicate in at least three independent assays.

### Invasion assays

Hep-2 cells were grown on sterile coverslips in MEM medium supplemented with 8% SFB for 24h until the monolayer reach 80% confluence. Tachyzoites were pretreated with the same conditions that for viability assay. Then Hep-2 cells were exposed to tachyzoites pretreated, after 2h of interaction, samples were processed for optical microscopy analysis. Briefly, cells were fixed with 10% formaldehyde 30 min, permeabilized 5 min with 0.1% Triton-X100 and washed with 1X PBS. The samples were staining with haematoxylin – eosin solution (Merck Millipore KGaA, Darmstadt, Germany), washed with 50 % ethanol and slides were mounted with a PBS: Glycerol (1:1) solution. The invasion process was evaluated counting 300 total cells, we considered as invaded cells, every cell that presented at least one parasitophorous vacuole on their cytoplasm. Quantitative analysis was performed in an AxioObserve microscope (Carl Zeiss, Mexico). This assay was realized by triplicate in at least three independent assays.

### Induction of changes in tachyzoites morphology by DHEA

Extracellular tachyzoites treated with DHEA, S-P or DHEA / S-P at several concentrations, for 30 min or 2 h, were processed for Transmition Electron Microscopy (TEM). Briefly, tachyzoites were resuspended in 2.5 % glutaraldehyde in 1X PBS in gentle agitation for 1 h, washed with 1X PBS, fixed with 1 % OsO4 1 h, and contrasted with 1 % aqueous uranyl acetate 2 h. Samples were dehydrated in increasing concentrations of ethanol (50-100 %), then were embedded in crescents concentrations of LR White resin (London Resin, England, Electron Microscopy Sciences, USA) and polymerized at 4°C for 36 h under UV lamp. The samples were processed with a ultramicrotome, serial cut were performed of around 10 μm of thickness and mounted in a sample holder, the ultrastructural analysis was performed in a Transmission Electron Microscope JEM200CX 200KV (JEOL Co., Tokyo, Japan), image analysis was performed using the Digital Micrograph program (TM 3.7.0 for GSM 1.2 by the Gatan Software Team).

### 2D SDS-PAGE

Whole extract of, intact or treated parasites with DHEA (10 μM), S-P (800 μM) and DHEA / S-P (10 / 800 μM) for 30 minutes, was obtained by lysis in 2D sample buffer; extracts were centrifuged at 10 000 rpm, soluble fractions were quantified in a Nanodrop™ 2000 (Thermo Scientific) at 280 nm. 100 μg of whole extracts, contained in 125 μL of rehydratation buffer, were load on ImmobilineTM DryStrip pH 3-10, 7 cm strips (GE Healthcare).

After 16 h of passive rehydratation, isoelectric focus was performed in a Protean IEF Cell (Bio-Rad Laboratories, Firmware Version: 1.80) by the supplier specifications. Strips were equilibrated in an equilibrium buffer (6 M urea, 0.375 M Tris-HCl pH 8.8, 2% SDS, 20% glycerol) with 0.5% dithiothreitol (DTT) for 10 min, and then with equilibrium buffer with 2.5 % iodoacetamide (IAA) for 10 min; strips were loaded in polyacrylamide precast gels (Mini-PROTEAN®TGXTM Precast Gels 4-20%, Bio-Rad) and electrophoresis was performed at 100 V; then gel were stained with silver nitrate and scanned in a HP Scanjet G4050 scanner.

### *In silico* analysis of 2D SDS-PAGE

Images were analyzed in the PDQuest Advanced-8.0.1 program. We determined the spots number for each condition, and then we matched all gels using the control as reference, in order to obtain the matched spots between conditions, their relative expression, as well as, their isoelectric point (IP) and molecular weight (Mr). The spots that exhibited differential expression between treatments were identified by its molecular weight and isoelectric point using the TagIdent program of ExPASy portal (https://web.expasy.org/tagident/).

### Modelling, docking and molecular dynamics of the cytochrome b5 family heme/steroid binding domain-containing protein

Initial model generation was accomplished by using the cloned sequence for *Toxoplasma gondii* **PGRMC** and submitting it to Rosetta Homology modeling [45]. Resulting models clustered close together for the selection of the best model. However, since the template is a PGRMC1 protein in complex with a heme group (PDB ID 4×8Y) [46], these models were refined using UCSF Chimera-Modeller plugin [47-48]. Then, their quality was evaluated using Molprobity [49]. The highest quality model was selected to perform ligand docking. Blind docking was performed using Vina 1.1.2 on the LNS supercomputer. All ligands were obtained from the ZINC database and converted to PDBQT format using the GUI provided by Autodock Tools. The receptor was kept rigid during docking. Docking employed a grid of dimensions 40 × 40 x 40 with a 1 Å grid size. Exhaustiveness was always set to 1000. Analysis of the docking results was performed in UCSF Chimera. The results presented in Table are the best candidates selected from the consensus score the three best results.

### Statistical analysis

Statistical analysis was performed with a variance analysis (ANOVA) of two ways that allowed determined simultaneity the effect of two variables (treatment and exposition time) with the tukey comparation prove. We use the program GraphPad Prism 6, analysis was considered significative different when p < 0.05.

## Acknowledgements

Financial support: This Project was partially supported by Instituto Nacional de Cancerología (SMH). Angélica Luna Nophal was supported by a fellowship from Consejo Nacional de Ciencia y Tecnología (CONACyT) - Mexico No. 300434. Grant IN-209719 from Programa de Apoyo a Proyectos de Innovación Tecnológica (PAPIIT), Dirección General de Asuntos del Personal Académico (DGAPA), Universidad Nacional Autónoma de México (UNAM) and Grant FC2016-2125 from Fronteras en la Ciencia, Consejo Nacional de Ciencia y Tecnología (CONACYT), both to Jorge Morales-Montor. Grant # IT-200120 to Pedro Ostoa-Saloma and Grant # IA202919 to Karen E Nava-Castro, both are from PAPIIT, DGAPA, UNAM. Carmen T. Gómez de León is recipient of a Post-Doctoral fellowship from Grant FC2016-2125 from Fronteras en la Ciencia, Consejo Nacional de Ciencia y Tecnología (CONACYT). Authors acknowledge to Ricardo Hernández Ávila from the IIB, UNAM, for his available technical assistance.

## Data availability

The datasets generated and analyzed during the current study are available from the corresponding author on reasonable request.

## Author contributions

Saé Muñiz-Hernández: Methodology, conceptualization, project administration and writing.

Carmen T. Gómez-de León: Experimentation, data analysis and writing. Angélica Luna Nophal: Experimentation.

Lenin Domínguez-Ramírez: Experimentation, data analysis and writing.

Olga Araceli Patrón Soberano: Experimentation and data analysis Karen E Nava-Castro: Data analysis.

Pedro Ostoa-Saloma: 2D SDS-PAGE methodology and resources

Jorge Morales-Montor: Writing (review and editing), data analysis and resources.

